# Parkin R274W mutation affects muscle physiology via the PARIS-PGC-1α pathway

**DOI:** 10.1101/2023.10.26.564192

**Authors:** Martina Sevegnani, Adriano Lama, Francesco Girardi, Michael W. Hess, Maria Paulina Castelo, Irene Pichler, Stefano Biressi, Giovanni Piccoli

## Abstract

Recessive mutations in the Parkin gene (*PRKN*) are the most common cause of young-onset inherited parkinsonism. Parkin is a multifunctional E3 ubiquitin ligase that plays a variety of roles in the cell including the degradation of proteins and the maintenance of mitochondrial homeostasis, integrity, and biogenesis via the PARIS-PGC-1α pathway. In 2001 the R275W mutation in the *PRKN* gene was identified in two unrelated families with a multigenerational history of postural tremor, dystonia and parkinsonism. Drosophila models of Parkin R275W showed selective and progressive degeneration of dopaminergic neuronal clusters, mitochondrial abnormalities, and prominent climbing defects. In the Prkn mouse orthologue, the amino acid R274 corresponds to human R275. Here we described a muscle phenotype in R274W +/+ mice and discovered that a defective Parkin-PARIS-PGC-1α pathway may impact mitochondrial biogenesis and, eventually, myoblast proliferation and differentiation.

## Introduction

Recessive mutations in Parkin gene (*PRKN*) are the most common cause of young-onset inherited parkinsonism (Lesage et al., 2020). *PRKN* is located on chromosome 6q25.2-q27 and encodes a 52 kDa protein called Parkin, a multifunctional E3 ubiquitin ligase (Seirafi et al., 2015). Parkin is highly expressed in the brain, heart, and skeletal muscle, and it is present both in the cytosol and marginally in the cell nucleus (Kitada et al., 1998). Parkin modulates mitochondrial homeostasis, biogenesis, quality control and degradation (Harper et al., 2018). Indeed, Parkin can translocate to damaged mitochondria upon membrane depolarization and promote their autophagy-mediated degradation, a process called mitophagy, in a PINK1-dependent manner. Parkin is important not only for mitochondria elimination but is also involved in their biogenesis. Parkin ubiquitinates PARIS, a transcriptional repressor of PGC-1α, one of the key regulators of mitochondrial biogenesis and function. PARIS ubiquitination leads to its proteasomal degradation, allowing PGC-1α transcription and subsequent expression of downstream genes such as Nrf-1, Nrf-2, and TFAM, which increase cellular respiration, energy utilization and mitochondrial biogenesis (Zheng et al., 2017).

Loss-of-function mutations in Parkin mouse models cause PARIS accumulation and a reduction in the size and the number of mitochondria in ventral midbrain neurons (Stevens et al., 2015). In 2001, Farrer and colleagues identified a novel 924C>T point mutation in exon 7 of *PRKN* that leads to the substitution of arginine with tryptophan at position 275 (R275W) in the RING1 domain of the Parkin protein (Farrer et al., 2001). In 2012 Ruffmann and his group identified the first pure heterozygous R275W patient with clinical features typical of sporadic PD, like late age of onset (62 years), severe neuronal loss in the substantia nigra, and diffuse Lewy pathology (Ruffmann et al., 2012). Unlike other common *PRKN* mutations that result in the complete ablation of enzyme function, R275W Parkin retains a residual E3 ubiquitin ligase activity (Lee et al., 2010). Overexpression of human R275W Parkin in *Drosophila* causes selective and progressive degeneration of dopaminergic neuronal clusters, decreased Parkin solubility and prominent climbing defects in mutant flies compared to wild type. *Drosophila* mutants also showed marked fiber degeneration in the flight muscles, accompanied by severe and pleiomorphic mitochondrial pathologies (Wang et al., 2007). In the Prkn mouse orthologue, the amino acid R274 corresponds to human R275. We have recently characterized the Parkin R274W +/+ mice, which show age-related nigrostriatal degeneration and motor impairment (Regoni et al., 2023). Here, we described a muscle phenotype in R274W +/+ mice and discovered that a defective Parkin-PARIS-PGC-1α pathway may impact mitochondrial biogenesis and, eventually, myoblast proliferation and differentiation.

## Materials and methods

### Animals

The University of Trento and the National Ministry of Health approved all animal protocols (authorization #162/2022-PR). Experiments were designed to limit the number of mice to be sacrificed respecting the 3Rs rule and according to national guidelines. All the experiments were done in compliance with EU Directive 2010/63 and national law D.L. 26/2014. Parkin R274W+/+ mice were generated via CRISPR/Cas9 technology by Taconic Biosciences. Transgenic animals were crossed with C57BL/6N mice for more than 10 generations to produce R274W+/- and WT littermates. Transgenic animals are viable and fertile. Animals were kept in a normal light/dark cycle (12 hours light, 12 hours dark) with food and water *ad libitum*. At 6-, 12- and 18-months Parkin R274W+/+, R274W+/- and wild-type mice were evaluated for behavioural performance. Male and female mice were used in the same ratio. After the motor assessment, mice were sacrificed according to guidelines with CO_2_ and then we gathered the skeletal muscle tissue. Gastrocnemius and soleus muscle were frozen in isopentane placed on liquid nitrogen and kept at -80°C.

### Behavioural tests

Behavioral assessment was performed as previously described (Pischedda et al., 2021). Briefly, before proceeding to motor tests, mice were habituated to the presence of the operator by handling for 5 min per day for a minimum of 5 days prior to the first behavioral test. Both male and female mice (in equal ratio) were used in behavioral tests. We did not detect any sex effect in motor tests (supplementary figure 1).

#### Rotarod

Balance and coordination were measured using the rotarod apparatus from Ugo Basile (Biological Research Apparatus). The rotarod was set at fixed acceleration of 12 rpm, and the time needed to fall off the rotarod was recorded. Mice were habituated to the task for four consecutive days. The fourth trial of the last test day was evaluated for statistical analysis. A cut-off of 300 seconds was imposed.

#### *Hanging Wire Test* (four limbs)

Mice were placed on a metal grid. The grid was kept upside down and total hanging time was recorded. The stopwatch was stopped when the animals had lost grip with all four limbs. Three trials were performed for each mouse, and the best performance was used for statistical analysis. A cut-off of 300 seconds was imposed.

## Grip Strength Test (anterior limbs)

Strength in the anterior limbs was measured using the grip strength apparatus from Ugo Basile (Biological Research Apparatus). The maximum strength was set to 500g/f at a rate of 30gf/s. Mice were placed so that their anterior limbs could grasp a horizontal bar, and gently pulled by the tail with a direction parallel to the ground. Mean peak force of three consecutive trials with one minute rest in between was measured using the Ugo Basile DCA software. Mean peak force was then normalized on the animal weight, measured on the same day of the test.

### Hematoxylin & Eosin Staining (H&E)

Soleus muscles were cut using Thermo Scientific HM525 NX cryostat at a temperature of -21°C. 14 μm transversal sections were cut and placed on Polysine™ Adhesion Microscope Slides (Epredia). Slides were stored at -20°C until further analysis. Sections were defrosted and washed in deionized water. Glass slides were covered with Mayer’s Hematoxylin (abcam) for 4 minutes. After 2 minutes of washing in distilled water, bluing was achieved by dipping slices into a Differentiation Solution (70% EtOH; 0.3% HCl) and Scott’s Solution. Slides were washed in water for 2 minutes, then in 70%, and 90% alcohol. Eosin Y solution (abcam) was applied for about 30 seconds. Samples were washed in absolute alcohol for 1 minute. After being washed in xylene for 3 minutes, slices were mounted using DPX mounting medium (Sigma). Samples were visualized in brightfield illumination using optical microscope Zeiss Axio Imager M2 equipped with a colour camera and analysed for signal intensity using FIJI software.

### Fibroblast isolation and culture

Mouse fibroblasts were collected from female and male wild type and Parkin R274W mouse ears. The tissue was incubated with 70% ethanol for 5 minutes. After air-drying, the ears were cut into small pieces and incubated for 60 minutes at 37°C with a 1:1 mixture of collagenase II (Worthington) and dispase II (Gibco). Smashed tissue was put in a 70 μM cell strainer and ground using a syringe plunger. The cell suspension was centrifuged twice for 7 minutes at 580g at 4°C. The supernatant was discarded, and the pellet was resuspended and grown in DMEM with 20% FBS, 1% L-glutamine and 1% Pen-Strep. A complete medium change was performed after 48 hours, and after that was replaced every third day. Cells were incubated at 37°C in a humified 5% CO2 incubator. Cells were split 1:10 every 4 days with 0.25% trypsin and cryopreserved at need in 90% FBS – 10% DMSO.

### Myoblasts and myotubes isolation and culture

Female and male wild type and R274W+/+ mice were sacrificed at 3 months of age, and all the limbs muscles were collected and minced. Disaggregated tissue was incubated in Muscle Dissociation Buffer (MDB, 490 U/mL collagenase II in HamF10, 1% Pen-Strep, and 1% L-glutamine) for 40 minutes at 37°C under 70 rpm agitation. A wash was performed in ice cold Wash Media (WM, 1% Pen-Strep, 1% L-glutamine, 10% Horse Serum in HamF10) followed by centrifugation at 1600 rpm for 5 minutes at 4°C. After supernatant removal, dispase II (11 U/mL in PBS) and collagenase II (1960 U/mL in PBS) were added to the pellet and incubation was performed at 37°C under 70 rpm agitation for 20 minutes. Supernatant was further dissociated using 18’’ and 20’’ syringe needles and 70 μm Nylon strainer. After each step, a wash in WM was performed followed by centrifugation at 1600 rpm for 10 minutes at 4°C. Muscle satellite cells were isolated from the resulting pellet using MACS® (magnetic cells separation) technology by Miltenyi Biotec. Isolated cells were plated in 60mm collagen-coated (Collagen Sigma C3867) Petri dishes and grown in Growth Medium (GM, 2% Pen-Strep, 1% L-glutamine, 20% FBS, 2.5 ng/mL FGFβ in HamF10). The medium was changed every second day. Cells were incubated at 37°C in a humified 5% CO2 incubator. Cells were split at around 70% of confluency with 0.25% trypsin and cryopreserved at need in 90% FBS – 10% DMSO. To induce differentiation, cells were plated at 20000 cells/cm2 onto matrigel-coated plates in GM. After 24 hours, cells were placed in Differentiation Medium (DM, 2% Pen-Strep, 1% L-glutamine, 2% Horse Serum in DMEM) for 72 hours.

### Plasmids and Transfection

The pcDNA 3.1-Flag-PGC-1α plasmid was purchased through Addgene (#45501). The empty pcDNA 3.1+ vector was used as a transfection control. *E. coli* DH5alpha were transformed to amplify the plasmid. Mouse primary fibroblasts were transfected with Lipofectamine 2000 (Life Technologies) following manufacturer’s instructions. The medium was changed after 24 hours. Cells were processed after 48 hours.

### RNA extraction and quantitative real-time PCR (qRT-PCR)

Total RNA was isolated from murine fibroblasts and myoblasts using Total RNA Purification Kit (Norgen), according to the manufacturer’s protocol and as described in (Verrillo et al., 2021). After extraction, RNA concentration was quantified with the NanoDrop 2000C spectrophotometer (Thermo Fisher Scientific). After DNAse treatment (Thermo Fisher Scientific, according to manufacturer’s instructions), complementary DNA (cDNA) was generated using qRT SuperMix (Bimake). The cDNAs were used for quantitative PCR (qPCR) exploiting iTaq Universal SYBR® Green Supermix and CFX96 RealTime System (BioRad) for 40 cycles. The primers utilized are shown in the Table 1. All samples were triplicated, and transcripts levels were normalized for the housekeeping genes relative abundance (RPL27). Data shown were produced using Bio-Rad CFX Manager software and analyzed according to ddCt algorithm (Livak and Schmittgen, 2001).

**Table 1.**
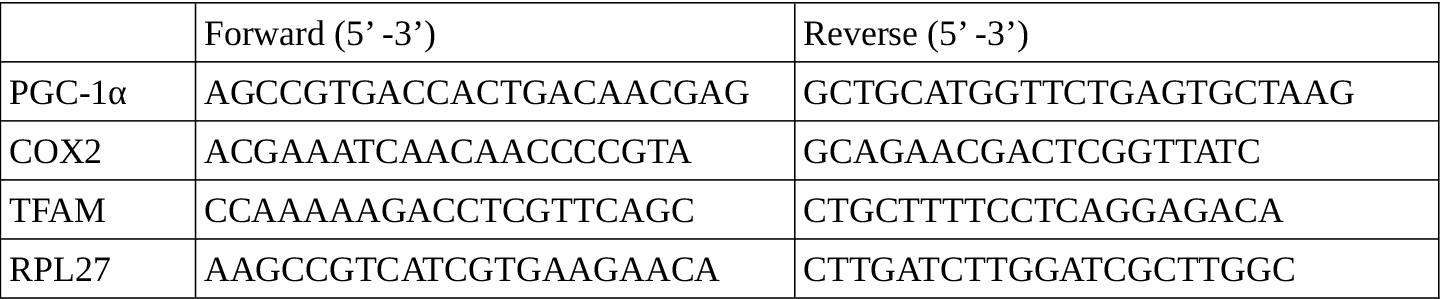
List of primers used in qRT-PCR assay.

### Western blotting

Western-blotting was performed as described in (Corti et al., 2008). Muscle tissues, murine fibroblasts and myoblasts were lysed in cold RIPA Buffer (150 mM NaCl, 2mM EDTA, 50mM Tris-HCl, 1% NP40, 0.25% sodium deoxycholate, pH 7.4) completed with protease inhibitors (Calbiochem) for 30 minutes at 4°C. After 10 minutes of 10000 g centrifugation at 4°C, Laemmli Buffer was added to the supernatant and samples were boiled for 10 minutes at 95°C. Skeletal muscle from WT or R274W mice were processed in the same way after the lysis in RIPA buffer using a 7 ml glass dounce tissue grinder (Wheaton). Samples were then loaded onto 10% or 15% handcast polyacrylamide gels. Proteins were transferred onto nitrocellulose membrane (Sigma-Aldrich) at 25V for 10 minutes using Trans-Blot Turbo Transfer (BioRad). After 1 hour saturation in blocking buffer (5% non-fat dry milk in 20 mM Tris, 150 mM NaCl, 0.1% Tween20, pH 7.4) primary antibodies were left overnight in blocking buffer at 4°C. Three washes of 10 minutes in TBS-Tween 0.1% were performed before 1 hour incubation with HRP-conjugated secondary antibodies in blocking buffer. The list of antibodies employed is presented in Table 2. ECL prime or select detection systems (Cynagen) were used to detect proteins, and images were acquired with the ChemiDoc Touch imaging system (BioRad). To obtain protein quantification, FIJI software was used to measure the optical density of the bands. For total protein analysis, β-actin was used as a housekeeping to normalize for protein content. Raw images are included in Supplementary figure 2.

**Table 2.** List of antibodies used in the study.

| Antigen | Host | Dilution | Code |
| --- | --- | --- | --- |
| Tomm20 | Rabbit | 1:500 | HPA011562 Sigma |
| PARIS | Rabbit | 1:500 | AB130867 Abcam |
| Parkin | Mouse | 1:500 | P6248 Sigma |
| β-actin | Mouse | 1:1000 | SC-811178 Santa Cruz |
| FLAG | Mouse | 1:1000 | SC-166355 Santa Cruz |

### Immunofluorescence

For cell cultures, cells were seeded at low density on ø12mm cover glasses (Thermo Fisher Scientific). Cells were fixed with PFA 4% for 10 minutes and washed with PBS. Brain sections were defrosted at RT and washed in PBS. Antigen retrieval was performed when needed using Citrate Buffer (sodium citrate 10 mM in dH2O, Tween 0.05%, pH 6) for 20 minutes at 100°C. After washing with PBS, samples were incubated for 1 hour in saturation buffer (2.5% BSA, 10% goat serum, 0.2% Triton). Subsequent washes were all performed with PBS-Triton 0.2%. Samples were incubated with primary antibodies in incubation buffer (2.5% BSA in PBS-Triton 0.2%) overnight at 4°C (table 2). After three washes of 10 minutes each, specimens were incubated with fluorescent secondary antibodies in incubation buffer for 1 hour. DAPI was used to visualize nuclei. After three washes of 10 minutes each, slides were mounted using FluorSaveTM Reagent (Millipore). Images were acquired using the optical microscope Zeiss Axio Imager M2 equipped with a 40X objective and analyzed for signal intensity using FIJI software.

### Electron Microscopy (EM)

Myoblasts were plated onto Thermanox plastic cell culture coverslips (Thermo Fisher Scientific) at 20000 cells/cm2 and differentiated into myotubes as already described. Samples were fixed with 2.5% glutaraldehyde (Sigma Aldrich) in 0.1 M phosphate buffer, followed by 0.5% aqueous OsO4 and plastic embedding. 100nm-thin sections were analysed by transmission EM..

### High-resolution respirometry

High-resolution respirometer (Oxygraph-2k, Oroboros Instruments) was used to measure mitochondrial respiration in primary mouse fibroblasts as described in (Castelo Rueda et al., 2023; Perez Carrion et al., 2018). First, 1×10^6^ cells were inserted in the chambers and permeabilized with digitonin (16.2 μM; Sigma) in MiR05 medium (10 mM KH2PO4, 60 mM lactobionic acid, 20 mM HEPES, 3 mM MgCl2, 0.5 mM EGTA, 20 mM taurine, 110 mM D-Sucrose and 1 mg/ml BSA fatty acid free). To characterize mitochondrial respiration, the following respiratory states were assessed: physiological respiration with endogenous substrates (routine respiration), complex I- and complex II-dependent respiration by addition of malate (M, 2mM), glutamate (G, 10 mM), ADP (D, 2.5 mM) and succinate (S, 10 mM), maximal capacity of the electron transfer system after stepwise titration of uncoupler (FCCP, 0.3 μM steps), respiration after complex I inhibition with Rotenone (Rot, 0.5 μM). Residual oxygen consumption (ROX) was measured after addition of antimycin A (Ama, 2.5 μM). Spare respiratory capacity was determined as the difference between maximal respiration and routine respiration. Absolute respiration values were normalized for the total number of cells (1×10^6^) and total activity of citrate synthase (nmol ml-1 min-1). Citrate Synthase was assayed spectrophotometrically at 412 nm and 30 °C on the EnVision 2105 Multimode Plate Reader (Perkin-Elmer), according to (Coore et al., 1971) with slight modifications. The final concentrations for the the reaction were acetyl-CoA 100 μM, oxalacetate 100 μM and 5,5′-dithiobis-(2-dinitrobenzoic acid) 100 μM in tris chloride buffer, pH 8 (100 mM). Following respirometry in the O2K, cells were removed from the chamber, and an aliquot was snap-frozen. Citrate synthase activity measurements were performed for each respirometry experiment, and averaged values were used for the normalization of respirometry results.All reagents were purchased from Sigma Aldrich.

### Statistical analysis

All data were expressed as the mean ± standard error of the mean (SEM). The normality of data distribution was determined using the D’Agostino and Pearson omnibus normality test, followed by an unpaired Student’s t test, ANOVA followed by Tuckey’s post-hoc test or two-way ANOVA followed by Bonferroni or Student’s t post-hoc test as appropriate on GraphPad PRISM ver. 5. The level of significance and the number of independent experiments (P, n) are indicated throughout the manuscript.

## Results

### The impact of Parkin R274W mutation on the skeletal muscle

The previous finding that R275W *Drosophila* model shows impairment in motor abilities linked to muscular strength (Wang et al., 2007) prompted us to evaluate whether skeletal muscle physiology may be affected in the R274W +/+ mouse model. To assess muscular capacity we tested wild type and homozygous mice at 6, 12, and 18 months on the rotarod at fixed acceleration (12 rpm), on the hanging wire and by the anterior limbs grip strength. While the performances were similar between genotypes at 6 months, we observed a decreased muscular performance in homozygous mice at 12 months in the rotarod at fixed acceleration [Figure 1D-F]. At 18 months, R274W+/+ mice performed poorly in all the tests performed [Figure 1G-I].

**Figure 1.**
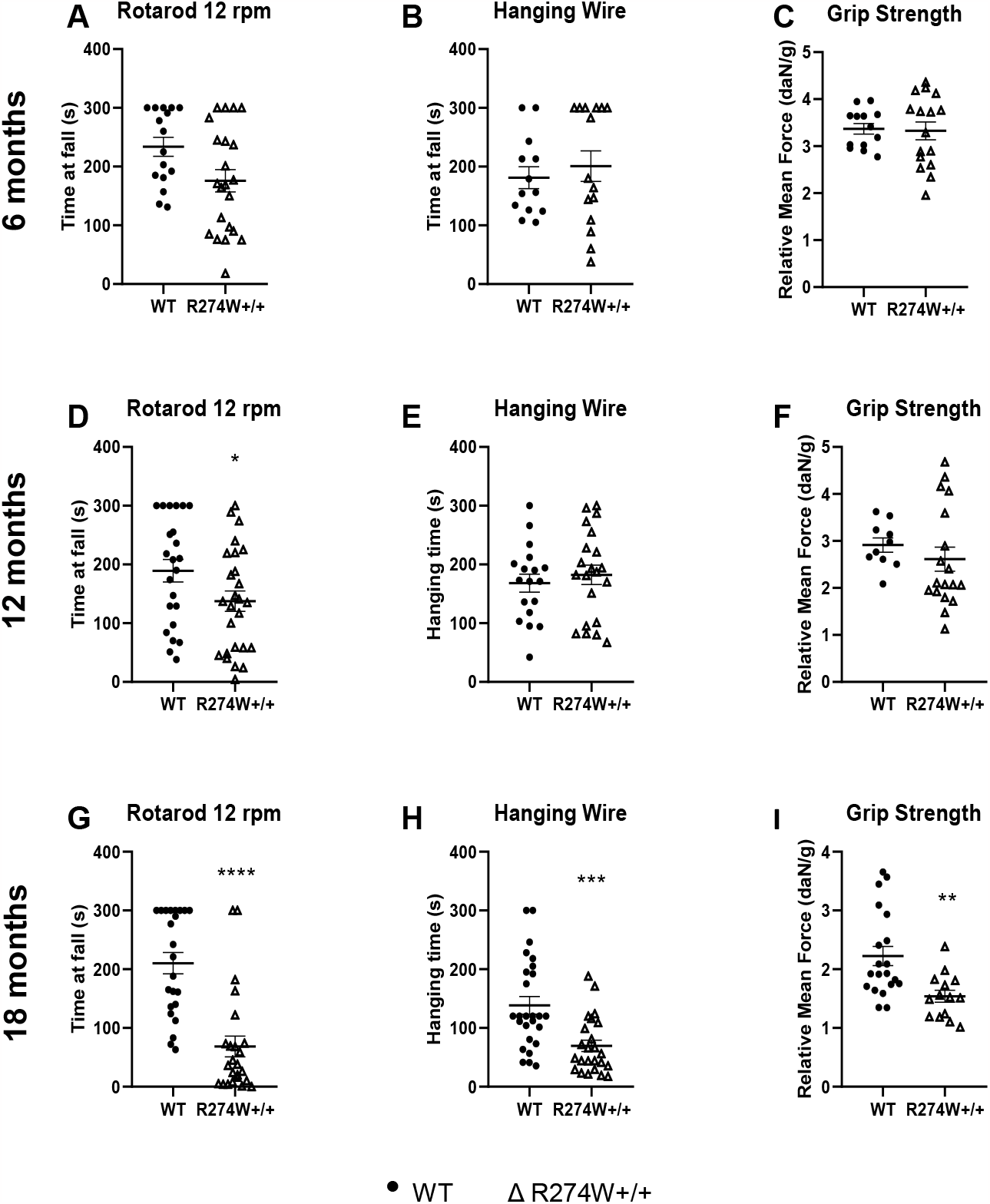
R274W+/+ mice show age-dependent impairment in muscular strength. We tested male and female mice at the indicated ages. We observed defects on the rotarod at fixed acceleration at 12 (D) and 18 months (G), and in the hanging wire and anterior limbs grip strength at 18 months (H, I). Data are the means ± SEM; n WT = 13-24, n R274W+/+ = 13-22; *, **, ***, ****, p<0.05, 0.01, 0.001, 0.0001 vs. wild type.

Altogether, these data show that R274W+/+ mice display an age-dependent strength impairment starting at 12 months of age. We have recently assessed that Parkin R274W is expressed at low level in mouse brain (Regoni et al., 2023). Accordingly, we detected by biochemical means a reduction in Parkin levels in 12-month-old homozygous soleus muscle compared to age-matched wild type [Figure 2A-B]. Next we analyzed skeletal muscle histology. We focused our analysis on homozygous mice at 12 and 18 months, when muscular performance is overtly impaired. Histological analyses were conducted on the soleus muscle, as it contains predominantly type I slow twitch fibers, the most involved in efforts of long duration (Carroll et al., 2011). Haematoxylin and eosin staining on transversal sections of the soleus muscle revealed a 20% and 26% decrease in the fibers cross-sectional area (CSA) of mutant mice compared to wild type respectively at 12 and 18 months, with an increase in the interstitial connective tissue that surrounds the muscle units [Figure 2C-E]. These findings suggest that homozygous mice present a degeneration in the skeletal muscle already at 12 months of age.

**Figure 2.**
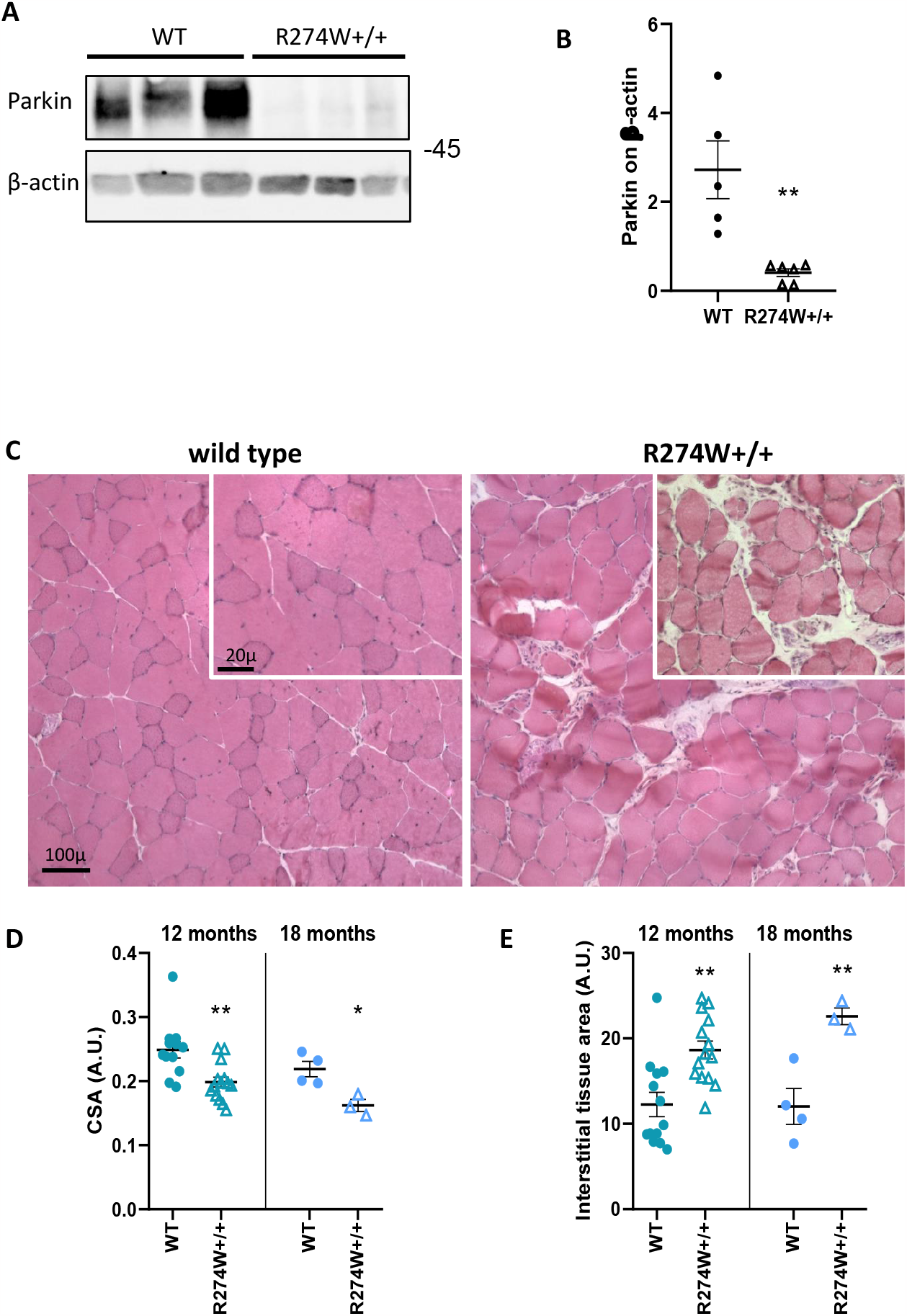
R274W mutation alters muscle tissue structure. (A) Parkin levels are reduced in 12-month-old homozygous mice compared to wild type in soleus muscle. (B) The graph shows Parkin amount normalized on β-actin. Data are expressed as means ±SEM; n = 5-6; **, p<0.01 versus wild type. (C) Representative images of haematoxylin and eosin staining on 14μm transversal sections of soleus muscle of 18-month-old wild type and homozygous mice. Decrease in muscle fiber size quantified as cross-sectional area (CSA) and increase in interstitial tissue are observed in homozygous tissue compared to wild type (D, E). Data are expressed as means ±SEM; n 12 months = 12-13, n 18 months = 4-3; *, **, p<0.05, 0.01.

### Parkin R274W +/+ derived muscle cells show alterations in proliferation, fusion ability, and mitochondrial defects

The reduced strength and defects in muscle structure observed in R274W +/+ mice may follow an impairment in muscle development. To this aim we derived myoblasts from wild type and homozygous mice. Myoblasts are precursors of muscle cells that differentiate *in vitro* into myocytes which eventually fuse together to form multinucleated myotubes. Myotubes are terminally differentiated cells useful for studying the characteristics of muscle fibers *in vitro* (Guo et al., 2020). The quantification of cell cycle kinetics by EdU (5-ethynyl-2′-deoxyuridine)-coupled-fluorescence-intensity analysis of myoblasts derived from wild type and homozygous mice showed that mutant cells are less proliferative, remaining mostly blocked in G1/G0 phase [Figure 3A-B]. Furthermore, myoblasts derived from homozygous mice showed a strong decrease in their fusion ability [Figure 4 A, B] and a 25% reduction in their diameter compared to wild type cells [Figure 4C]. These results suggest that R274W substitution in Parkin leads to defects in myoblasts differentiation and fusion. At the subcellular level, electron microscopy (EM) of 100nm-thin plastic sections showed normal ultrastructure of the mitochondrial network in WT myotubes. Throughout the entire cell population mitochondria presented as elongated ∼220nm wide rods with local, moderate narrowings and occasional branches. R274 +/+ cells, however, were characterized by a high diversity of partly abnormally shaped mitochondria. Apart from a few, virtually “normal”, branching rods, there regularly occurred comparatively huge, pleomorphic organelle profiles (up to >800nm in diameter), sometimes forming only ∼120nm-wide, rod-shaped protrusions. Similar slim rods displaying envelope and cristae (i.e., mitochondrial characteristics) appeared to lie freely within the cytoplasm, but this phenomenon might result from adverse section planes and/or artifactual organelle fragmentation during specimen fixation [Figure 5A]. It is well established that Parkin controls mitochondrial biogenesis via the modulation of PARIS degradation. PARIS represses the transcription of PGC-1α, a master regulator of mitochondrial biogenesis. Parkin LOF induces PARIS accumulation, down-regulates PGC-1α transcription, and eventually leads to decreased mitochondrial biogenesis. By biochemical measn, we found that parkin protein is down-regulated in R274W+/+ myoblasts. We also observed an increase of PARIS and a reduction of the mitochondrial marker Tomm20 in homozygous myoblasts compared to wild type [Figure 5C-F]. In addition, we observed that the mRNA of PGC-1α and its target TFAM, a crucial actor in mitochondrial biogenesis, as well as of COX2, a mtDNA-encoded gene [Figure 5 G-I] are reduced in mutant cells. Altogether, our data indicate a reduction in mitochondrial biogenesis in R274W +/+ cells.

**Figure 3.**
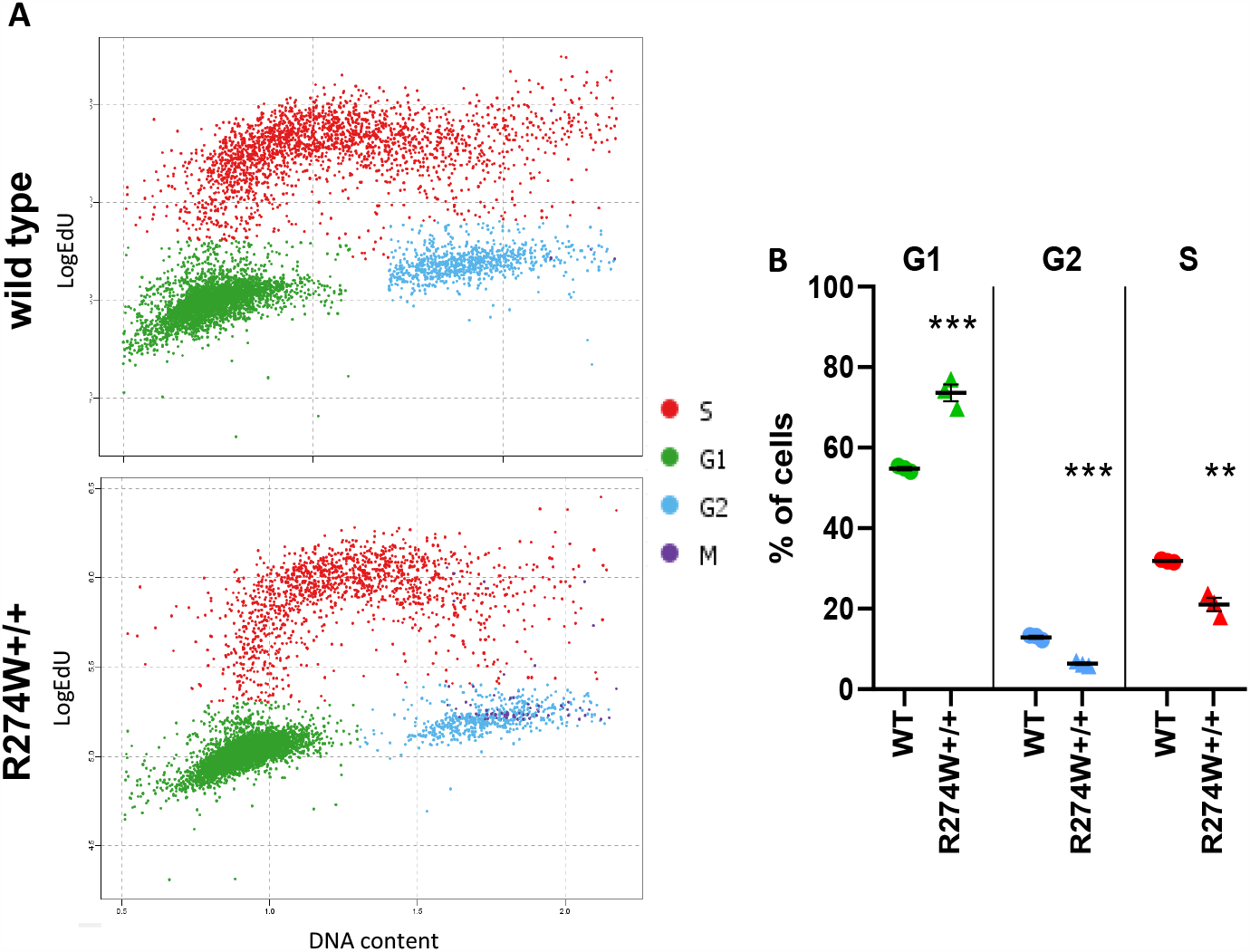
Parkin R274W+/+ affects myoblasts proliferation. We analysed proliferative behaviour of wild type and homozygous myoblasts by means of a EdU assay. Only proliferative cells (G2, S phase) are able to incorporate fluorescently labelled EdU. Phospho-histone H3 (PH3) was used to specifically stain cells undergoing mitosis (M). DNA content (DAPI) was used to normalize the levels of fluorescent markers. (A) Each point of the graph represents a single cell. The relative position on x-(DNA content) and y-axis (LogEdU) was used to distinguish between different cell cycle phases. (B) The graph shows percentage of cells in G1/G0, G2, and M phase. Each point represents the mean values of all single cells for each well analysed. Data are expressed as means ±SEM, n = 3; **, ***, p<0.01, 0.001, versus wild type.

**Figure 4.**
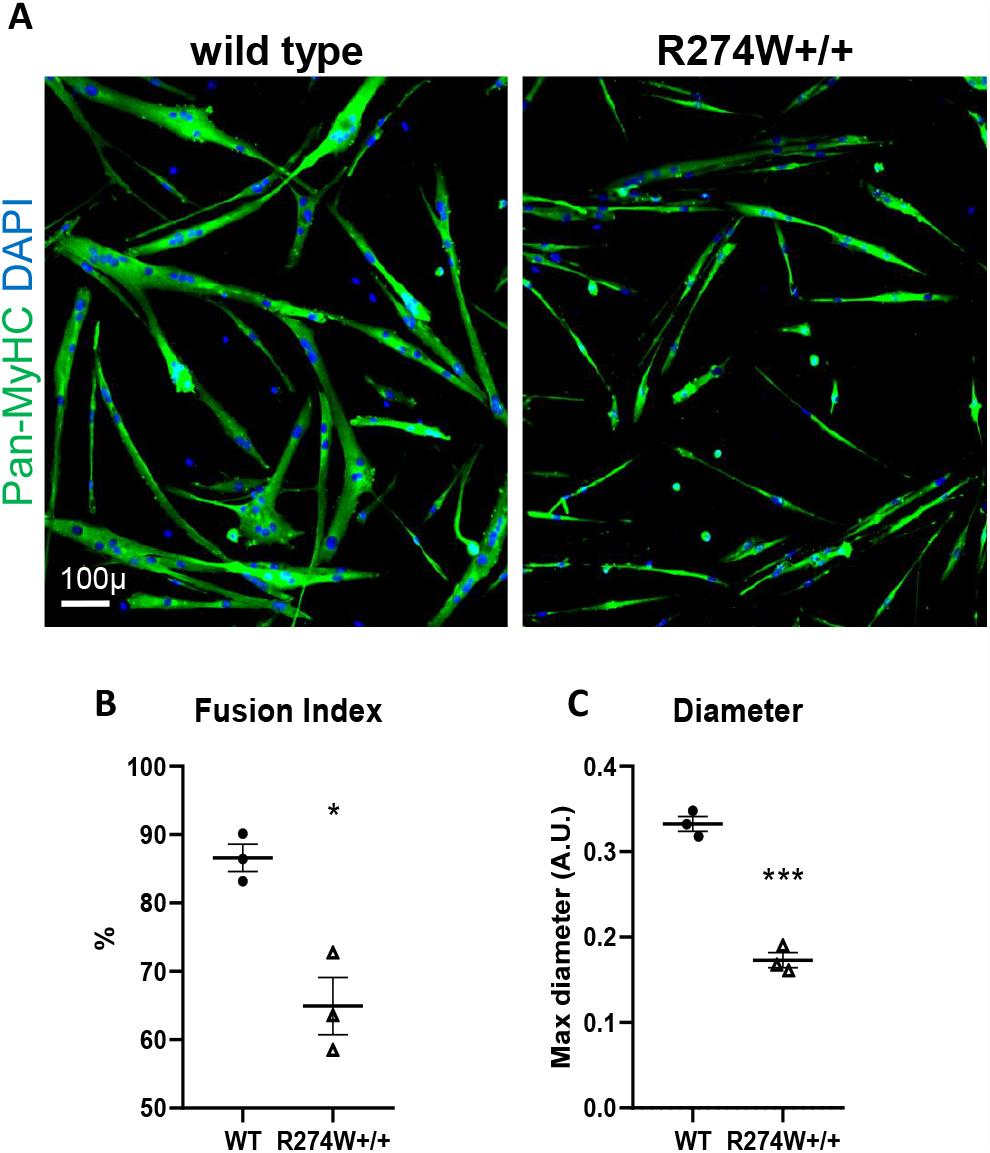
R274W Parkin impairs myotubes formation. (A) We differentiated myotubes from wild type and R274W+/+ myoblasts. Myogenic differentiation was evaluated by the expression of myosin heavy chain using Pan-MyHC antibodies (green). Nuclei are visualized by DAPI in blue. Scale bar = 100μm. Fusion index (B) was calculated as the number of multinucleated cells folded over total number of myosin-expressing differentiated cells (green). (C) The graphs show maximum diameter of wild type and homozygous myotubes. In graphs (B, C) each point represents the mean of three technical replicates of a single cell line derived from a different mouse. Data are expressed as means ±SEM, n = 3; *, ***, p<0.05, 0.01, versus wild type.

**Figure 5.**
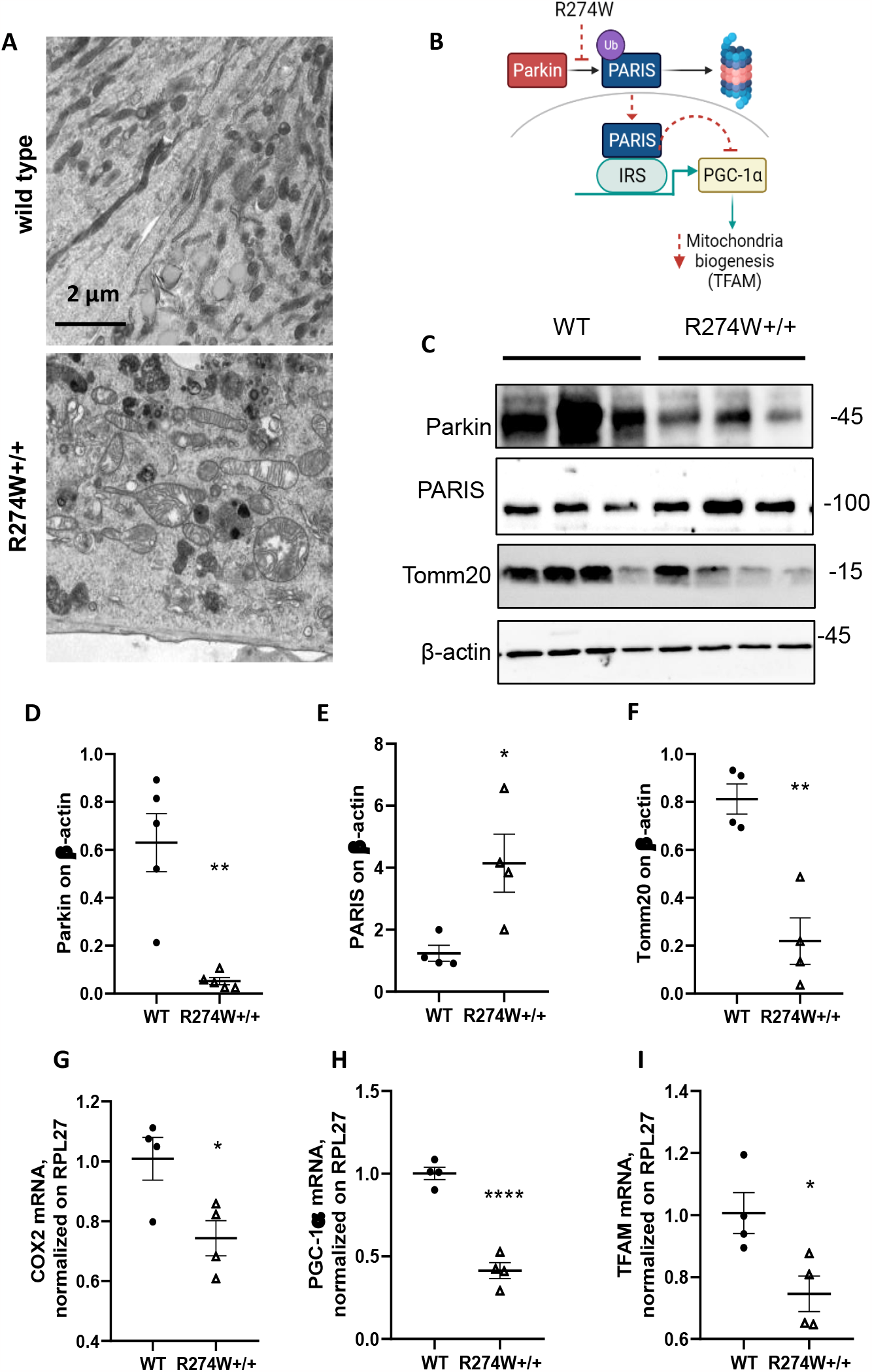
R274W+/+ myotubes show mitochondrial defects. (A) EM reveals normal mitochondrial networks (asterisks) in wild type myotubes. R274W+/+ myotubes show enlarged, pleomorphic mitochondria (asterisks) with slim protrusions (arrows) and/or virtually isolated, slim mitochondrial “rods” (arrows) nearby. Scale bar = 2μm. (B) Schematic overview of Parkin-PARIS-GC-1α pathway. Parkin controls PARIS degradation via the proteasome. PARIS represses the transcription of PGC-1α binding the insulin response sequence (IRS) region of its promoter. PGC-1α is a master regulator of mitochondrial biogenesis as it promotes the transcription of key genes such as the mitochondrial transcription factor A (TFAM). Parkin loss-of-function induces PARIS accumulation, down-regulates PGC-1α transcription, and eventually leads to decreased mitochondrial biogenesis. (C) We analysed wild type and R274W+/+ myotubes for Parkin, PARIS, and Tomm20 content via western blot. The graphs (D-F) show proteins level normalized on β-actin. Data are expressed as means ±SEM; n = 5; **, p<0.01 versus wild type. (G-I) The graphs show mRNA levels of COX2, PGC-1α, and TFAM normalized on RPL27 mRNA. Results are reported as means ±SEM, n = 4; *, ****, p<0.05, 0.0001, versus wild type.

### PARIS-PGC-1α pathway is altered in Parkin R274W +/+ fibroblasts

We validated the impact of Parkin R274W on the PARIS-PGC-1α pathway in primary fibroblasts generated from wild type and homozygous mice. First, we observed by western blotting that Parkin and Tomm20 protein levels were reduced in R274W+/+ fibroblasts compared to wild type cells [Figure 6B-C]. In mutant fibroblasts we also noticed the accumulation of PARIS protein [Figure 6D] and the reduction of PGC-1α mRNA [Figure 6E]. By high resolution respirometry, we observed a general decrease in the respiratory capacity, detected in the Routine state, complex I and II-dependent respiration, the maximal respiratory capacity as well as the respiration after complex I inhibition (Rot) in homozygous cells compared to wild type cells. Spare respiratory capacity indicates the mitochondrial capability to respond to a chronic mitochondrial insult, which might be the presence of a Parkin mutation, and this parameter was also significantly reduced in the mutant cells [Figure 6F]. Finally, we assessed mitochondrial content by imaging and measured a reduction in mitochondrial content in R274W +/+ cell. Interestingly, the overexpression of PGC-1α is sufficient to restore the mitochondria content in mutant fibroblasts [Figure 6 G-H]. Our data may suggest that PARIS upregulation impairs PGC-1α dependent mitochondrial biogenesis in R274W +/+ cells.

**Figure 6.**
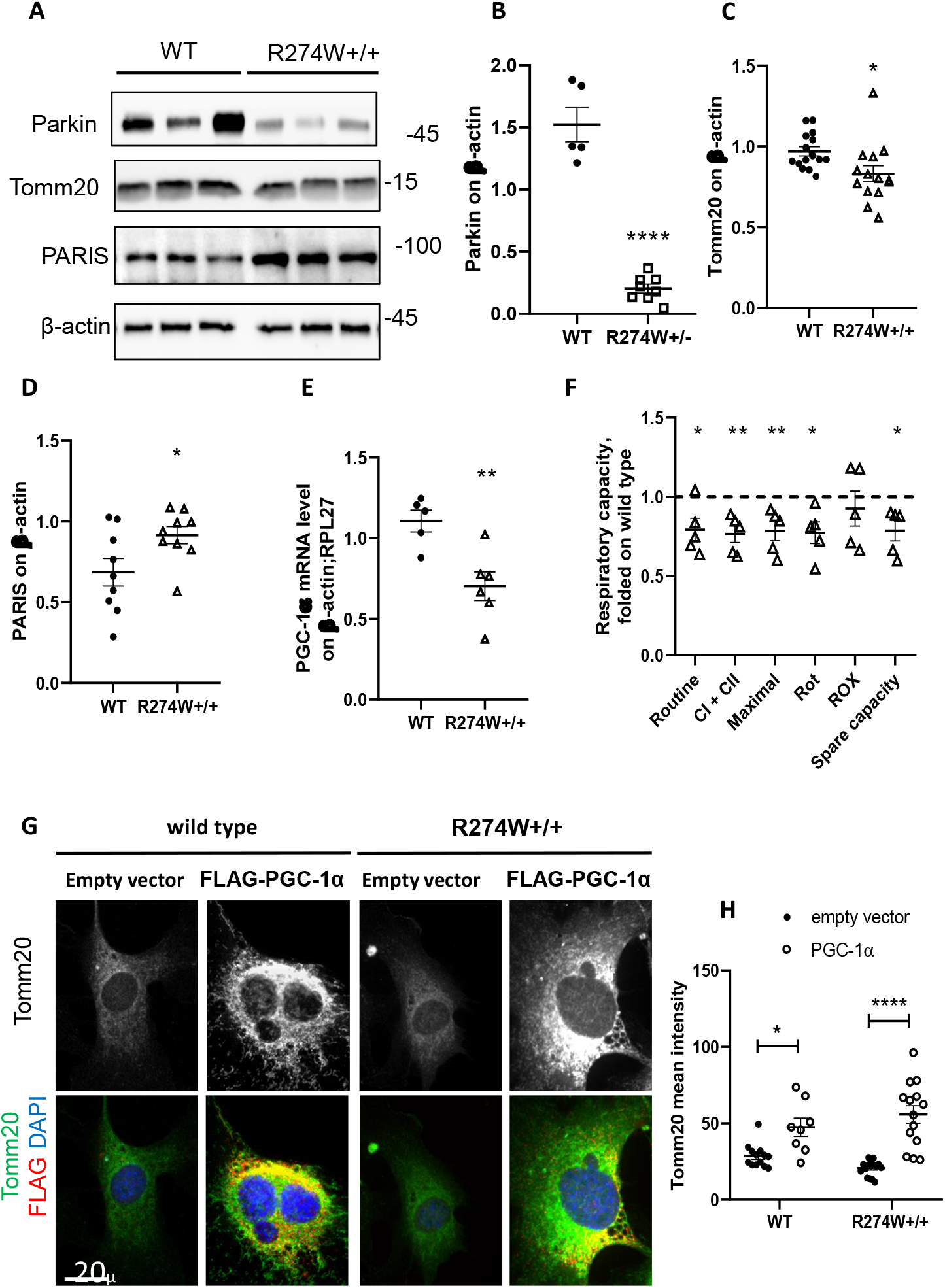
The PARIS-PGC-1α pathway is impaired in R274W+/+ fibroblasts. (A) We analysed by western-blotting samples prepared from wild type and Parkin R274W+/+ primary fibroblasts. The graphs report Tomm20 (B) and PARIS (C) levels expressed as optical density and normalized on β-actin amount. Data are expressed as means ±SEM, (B) n = 15-14; (C) n = 9; *, p<0.05 versus wild type. (E) We assessed PGC-1α mRNA level in wild type and Parkin R274W+/+ primary fibroblasts by qRT-PCR. The graph represents the relative expression of PGC-1α mRNA, normalized on β-actin and RPL27 mRNA amount. Data are expressed as means ±SEM, n = 5-6; **, p<0.01 versus wild type. (F) We assessed oxygen consumption in wild type and homozygous primary fibroblasts by high resolution respirometry. The graph reports routine respiration, complex I and complex II-dependent respiration (upon treatment with malate, glutamate, ADP and succinate), maximal respiration (maximal uncoupled respiration after uncoupler addition), respiration after inhibition of complex I (upon Rotenone addition), residual oxygen consumption (ROX) due to oxidative side reactions (after addition of antimycin A), and difference between maximal respiration and routine respiration (spare respiratory capacity). Data are expressed as means ±SEM; n = 5; *, **, p<0.05, 0.01 versus wild type. (G) We transfected primary fibroblasts derived from wild type and Parkin R274W+/+ mice with empty vector or with FLAG-PGC-1α vector. Cells were stained with anti-FLAG (red) and anti-Tomm20 (green) antibodies and DAPI to visualize the nuclei. Scale bar = 20 μm. The graph (H) shows that overexpression of PGC-1α increases Tomm20 levels. Data are expressed as means ±SEM; n = 8; *, ****, p<0.05, 0.0001 versus wild type.

## Discussion

Beside the well-known symptomatology related to tremor, gait, and postural imbalance, PD in humans is characterized by motor symptoms that are not strictly correlated with dopamine depletion in the extrapyramidal system. For example, bilateral muscle weakness and fatigue are often present in PD patients with unilateral parkinsonism (Cano-de-la-Cuerda et al., 2010). It is still not clear whether impairments in the muscle are caused by central deficits due to lack of dopamine (Frazzitta et al., 2015), incorrect muscular stimulation and stiffness, or other peripheral causes secondarily linked to PD like ATP depletion, mitochondrial degeneration, and inflammation (Kostić et al., 2016). Parkin R274W homozygous mice display defects in muscular strength and increased fatigue compared to wild type mice starting from 12 months of age. Accordingly, our *ex vivo* analysis of muscle tissue revealed alterations in the structure and organization of the muscle fibers, with decreased diameter and enlargement of interstitial tissue, similar to observation on R275W *Drosophila* (Wang et al., 2007). Coherent with these data, EM analysis of myotubes derived from our R274W+/+ mice regularly showed seriously damaged mitochondria. Most conspicuous were their enlarged, pleomorphic shape, their slim protrusions and virtually isolated slim mitochondrial “rods”nearby. Additionally, muscle cells derived from R274W+/+ mice also displayed alterations in their ability to replicate and fuse, processes that deeply rely on the correct functionality of the mitochondrial respiratory chain. In fact, myoblasts differentiation and fusion processes strongly rely on ATP levels (O’Connor et al., 2008). To obtain insight into the mechanisms that link Parkin R274W mutation, impaired mitochondria, and defects in muscle tissue we explored the PARIS-PGC-1α pathway of mitochondrial biogenesis, which is modulated by Parkin (Shin et al., 2011). PGC-1α is recognized as one of the major regulator of mitochondrial content and oxidative metabolism in several tissues, including skeletal muscle (Handschin and Spiegelman, 2006). We noticed that PARIS accumulates in homozygous muscle cells.

PARIS accumulation has been put in relation to ROS production that may damage mitochondria and reduce cell proliferation (Bae et al., 2020). On the other hand, the subsequent downregulation of PGC-1α and its downstream target genes may lead to the reduction in mitochondrial mass that we observed in homozygous myotubes and fibroblasts both by qRT-PCR and biochemical assays. Altogether, these data show that PARIS upregulation due to Parkin R274W mutation impairs mitochondrial biogenesis, reduces oxidative metabolism, and eventually affects myoblast proliferation and fusion. *In vivo*, this pathological cascade may result in reduced muscle fiber diameter and therefore, muscle strength.

We observed a significant deterioration of muscle strength only in 12- and 18-months mice, suggesting the occurrence of an age related mechanism. The life expectancy of mice is approximately 24 months. Muscular strength, exercise endurance, muscle volume, and muscle mass are significantly impaired in 18-month-old mice, suggesting the presence of sarcopenia (Kim and Hwang, 2020). Aging is one of the main contributing factors for most musculoskeletal diseases, including sarcopenia (Sharma et al., 2023). Ageing is characterized by an increased generation of ROS (Calvani et al., 2013) and lower gene and protein expression of PGC-1α (Joseph et al., 2012). An impaired mitochondrial function and reduced biogenesis may contribute to muscle atrophy observed in ageing (Kubat et al., 2023). The PARIS-PGC-1α axis is reportedly involved in different aspects of skeletal muscle biology, which include the regeneration upon damage, the adaptive response to exercise, the control of fiber typing, and the control of mitochondrial dynamics (Adhihetty et al., 2009; Dinulovic et al., 2016b; Lin et al., 2002). The alteration in PARIS-PGC-1α pathway and the reduction in mitochondrial content observed in Parkin R274W models may eventually exacerbate the age-related sarcopenia.

It is well established that adult skeletal muscle possesses a remarkable capacity for regeneration endowed by a population of adult resident stem cells, the satellite cells (Relaix and Zammit, 2012). Increasing observations suggest an association between muscle atrophy and satellite cell dysfunction (Biressi and Gopinath, 2015). Particularly, the loss of satellite cells is reportedly associated with age-related skeletal muscle atrophy in mice (Liu et al., 2017). A large body of evidence indicates that intrinsic and extrinsic satellite cell defects are mediating an impaired regeneration in old muscle (Muñoz-Cánoves et al., 2020).

These defects include metabolic and mitochondrial dysfunction (García-Prat et al., 2016). The PARIS-PGC-1α axis controls the lineage progression of satellite cells (Dinulovic et al., 2016a) Given the severe proliferative and differentiation dysfunction that we report here for satellite cell-derived Parkin R274W myoblasts, it is tempting to speculate that the Parkin mutation might affect via the PARIS-PGC-1α pathway the satellite cell population and, eventually, accelerate age-related sarcopenia in Parkin R274W mice.

## Conclusions

The correct functionality of Parkin is important for mitochondrial biogenesis and tissue functionality not only in the CNS but also in the periphery. The modulation of the PARIS-PGC-1α axis, including downstream target as Nrf-1, Nrf-2, and TFAM may represent a therapeutic target to ameliorate skeletal muscle performances in Parkin-PD patients. Intriguingly, a recent report identified farnesol as a potent inducer of PGC-1α, capable of ameliorating aging-associated muscle wasting through enhanced PARIS farnesylation, thereby relieving PARIS-mediated PGC-1α suppression (Bae et al., 2023). This observation calls for further investigation of this drug in PD models. A clear limitation of this study is that we mainly focused on the role of Parkin in mitochondria biogenesis. Future work will consider the effect of R274W mutation on Parkin-related mitophagy and its impact on muscle physiology. In fact, Parkin -/- mice demonstrated inefficient mitophagy and increased mitochondrial ROS, resulting in the acceleration of skeletal muscle atrophy (Chen et al., 2018a, 2018b; Ito et al., 2022).

## Author Contributions

conceptualization M.S. and G.P.; methodology M.S., A.L., MP.C and F.G.; data curation M.S., I.P, M.H, S.B. and G.P.; writing M.S, S.B., and G.P.; funding acquisition G.P.. All authors have read and agreed to the published version of the manuscript.

## Institutional Review Board Statement

Animal care and experimental procedures were conducted in accordance with the University of Trento Ethics committee and were approved by the Italian Ministry of Health, following the guidelines of the European Union, approved by the Italian Ministry of Health.

## Data Availability Statement

the full data set is available upon reasonable request.

