## Supplementary figures and images for "Parkin R274W mutation affects muscle physiology via the PARIS-PGC-1α pathway"

### supplementary figure 1

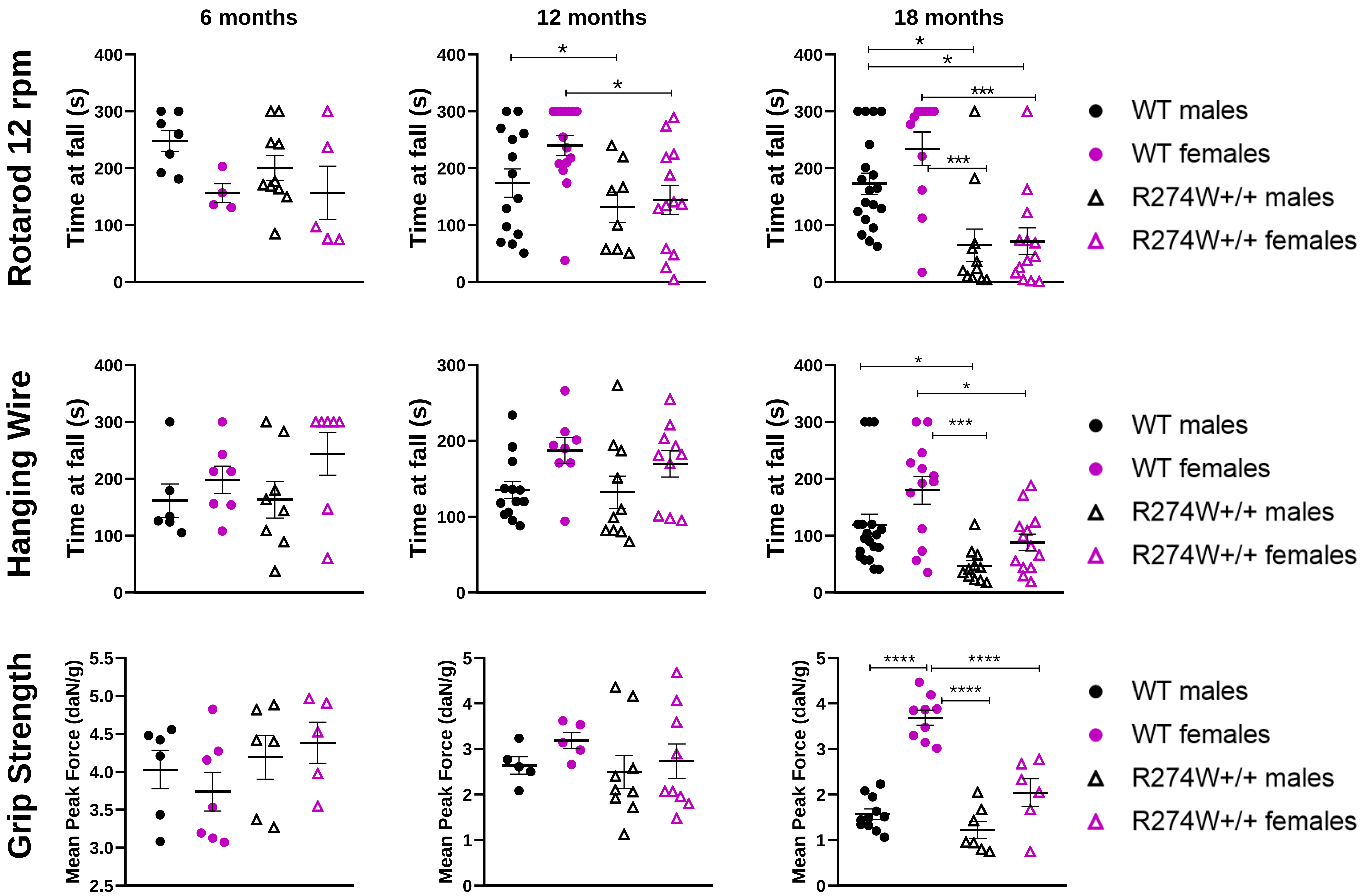
