## supplementary figure 2 for "Parkin R274W mutation affects muscle physiology via the PARIS-PGC-1α pathway"

Figure 2A: anti Parkin

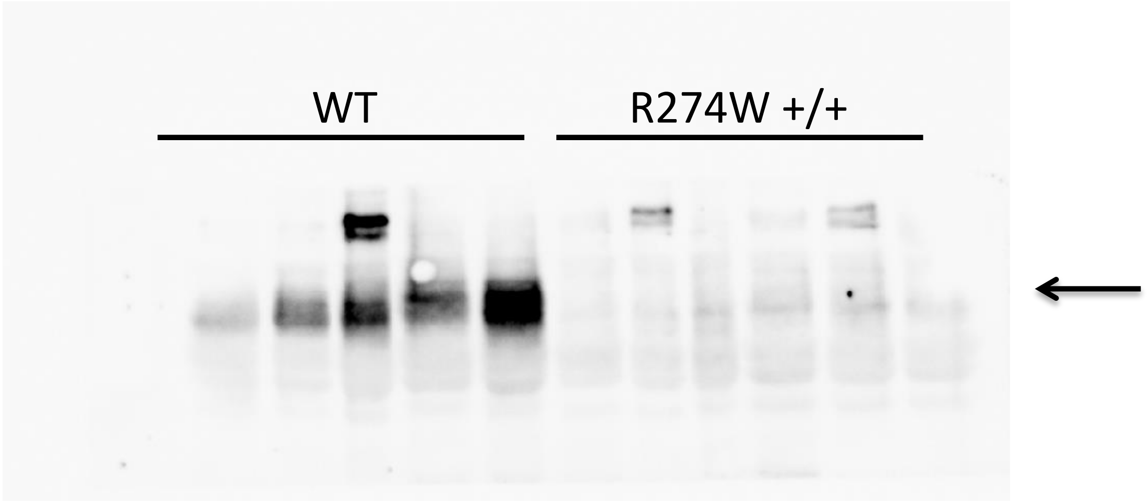


Figure 2A: anti b actin

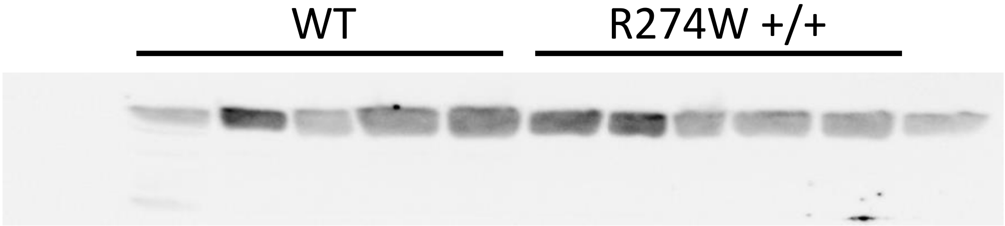

Figure 5C: anti Parkin

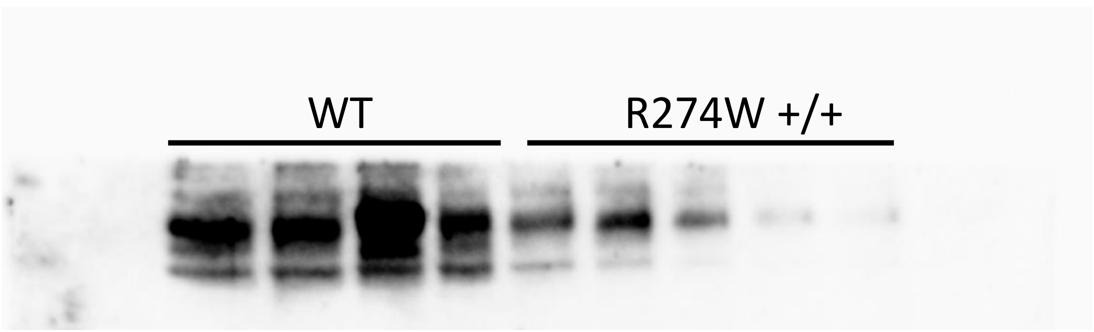


Figure 5C: anti Paris

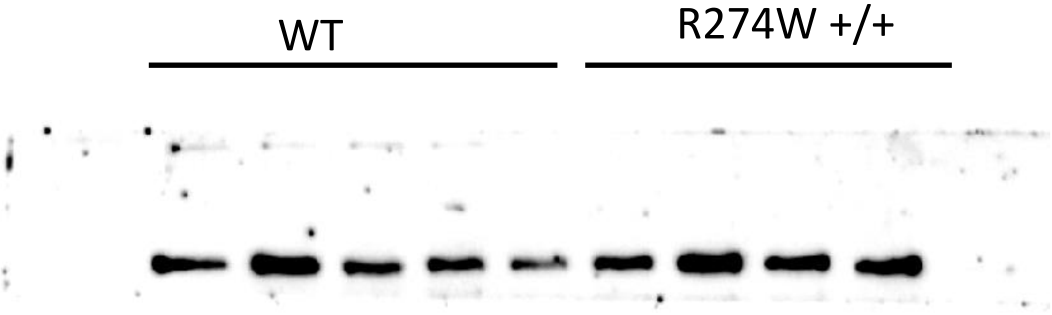


Figure 5C: anti Tomm20

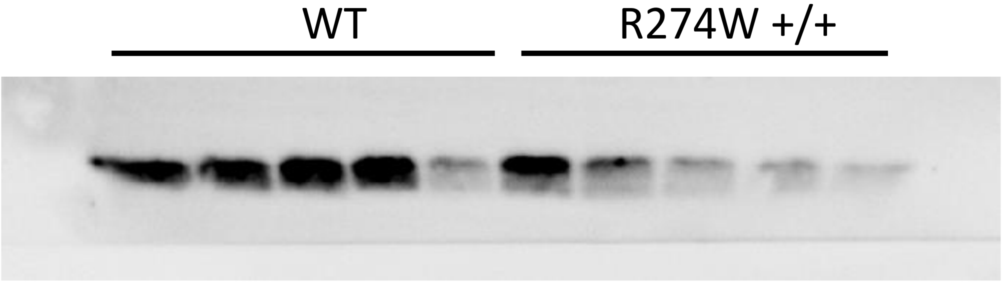


Figure 5C: anti b actin

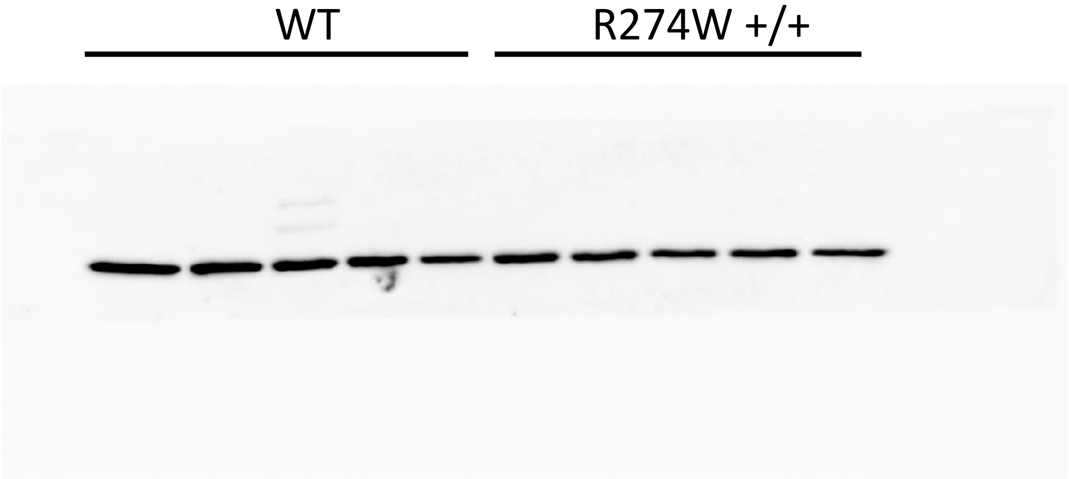

Figure 6A: anti Parkin

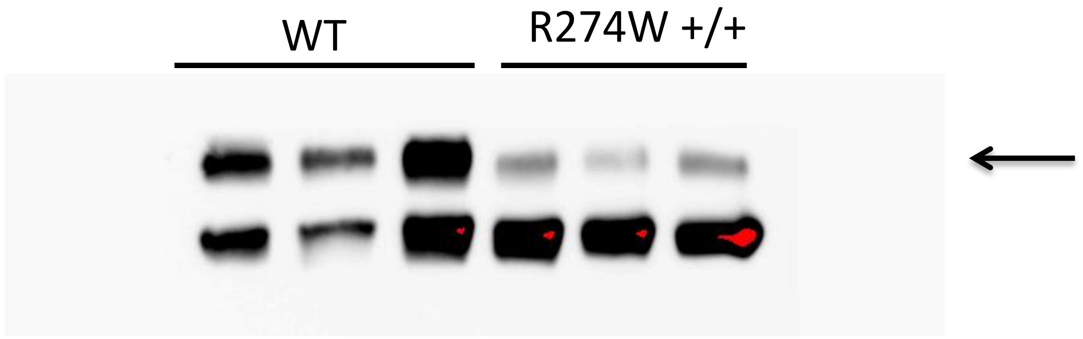


Figure 6A: anti Tomm20

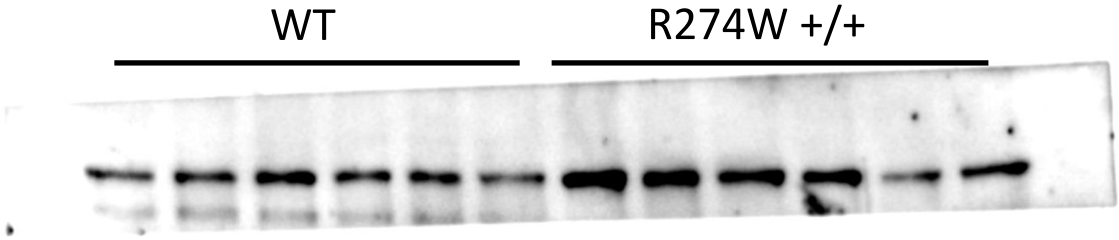


Figure 6A: anti Tomm20

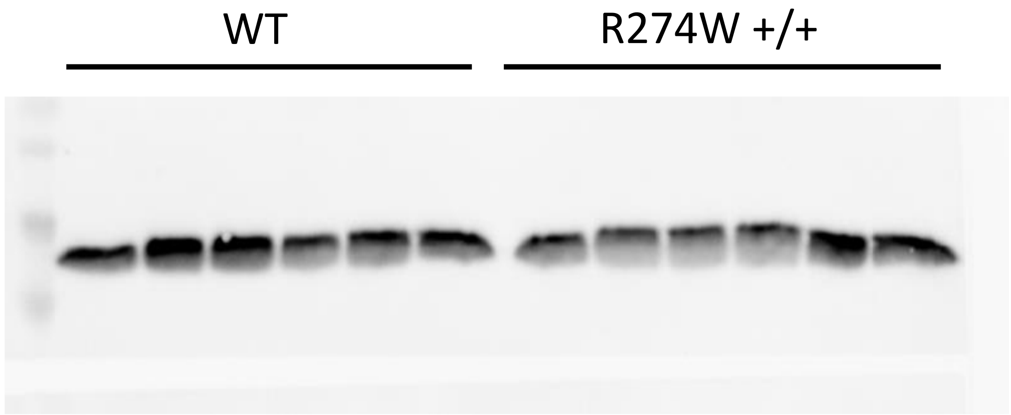


Figure 6A: anti Tomm20

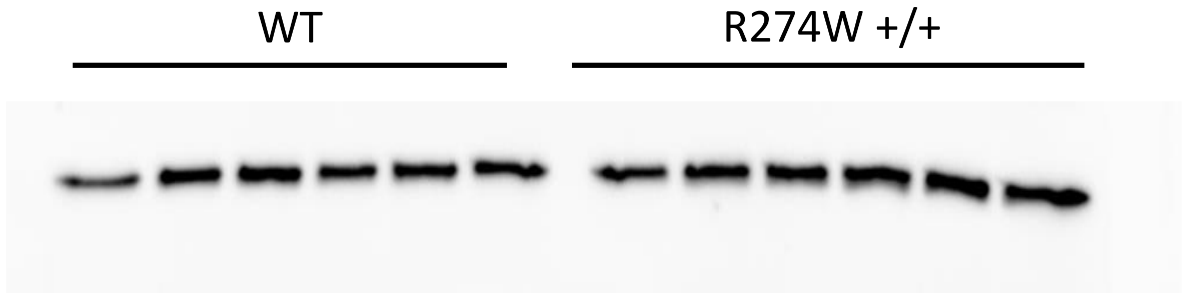
